# Modelling collective navigation via nonlocal communication

**DOI:** 10.1101/2021.05.09.443340

**Authors:** S. T. Johnston, K. J. Painter

## Abstract

Collective migration occurs throughout the animal kingdom, and demands both the interpretation of navigational cues and the perception of other individuals within the group. Navigational cues orient individuals toward a destination, while it has been demonstrated that communication between individuals enhances navigation through a reduction in orientation error. We develop a mathematical model of collective navigation that synthesises navigational cues and perception of other individuals. Crucially, this approach incorporates the uncertainty inherent to cue interpretation and perception in the decision making process, which can arise due to noisy environments. We demonstrate that collective navigation is more efficient than individual navigation, provided a threshold number of other individuals are perceptible. This benefit is even more pronounced in low navigation information environments. In navigation “blindspots”, where no information is available, navigation is enhanced through a relay that connects individuals in information-poor regions to individuals in information-rich regions. As an expository case study, we apply our framework to minke whale migration in the North East Atlantic Ocean, and quantify the decrease in navigation ability due to anthropogenic noise pollution.

## Introduction

Many animals routinely migrate long distances; spectacular examples include the pole-to-pole flights of Arctic terns and the transoceanic migrations of many whale species [1, 2]. Topography, the geomagnetic field, celestial information, and chemical signals can all serve to orient animals en-route [3, 4, 5]. Navigational cues may be interpreted in combination, or an animal may switch from a cue suitable for long-distance migration to a cue suitable for precise navigation when close to the destination [6, 7]. Frequently, migrations are conducted as a group and there is significant interest in the extent to which the “wisdom of the crowd” improves navigation performance. Improvement may arise from group heterogeneity, where knowledgeable individuals take on a leadership role, but is also hypothesised to occur in a homogeneous population through the “many wrongs” principal of navigation. Here, pooling information across the group reduces individual-level uncertainty via averaging[8, 9, 10, 11]. As an example, homing pigeons display improved homing behaviour when travelling in a small flock, compared to when flying solo [12].

Collective navigation demands communication with, or perception of, other group members. These interactions in turn influence an individual’s behavioural response [13, 14]. The complexity and range of interaction will vary significantly with an animal’s sensory machinery, along with the environment through which the animals are moving. For example, sound transmission through water permits whales to communicate with each other through “whalesong” and other vocalisations, up to estimated distances of hundreds of kilometres [15, 7]. Even on land, calls may travel several kilometres between elephants [16]. As such, a superficially dispersed animal population may be migrating as a group through communication across long distances.

Navigation also requires a robust evaluation of orienteering cues; the quality of orienteering information is unlikely to be uniform across the travel route. In animal populations with established social networks, such as whale societies, information from a dominant member may be considered more valuable than information from other members [17, 18], suggesting that the navigation may be easier in the presence of knowledgeable individuals. Alternatively, as the distance between the navigating individual and its target decreases, cues may become stronger, as for audible information, or weaker, as for detecting geomagnetic field differences. Journeys may require passage through *blindspots*, regions of space with diminished quality of navigational information. Blindspots may form naturally, for example due to adverse weather conditions, or through anthropological activity. Recently, significant attention has been devoted to the state of the marine “soundscape” [19]. Human oceanic activity has substantially increased over the last century, with extreme noise sources and raised ambient noise levels a result of shipping, offshore construction, and naval operations [20, 21]. This anthropogenic noise pollution has had a broad impact on the ocean-dwelling organisms that rely on auditory information [19]. For example, various cetaceans adjust the volume and frequency of their calls to account for marine noise [22, 23, 24], a behaviour known as the Lombard effect. However, this only provides partial compensation and the adjusted calls may encode less information. It has been estimated that species including minke whales (*Balaenoptera acutorostrata*) and humpback whales (*Megaptera novaeangliae*) could lose *∼*80% of their communication distance in the presence of increased human activity [22, 23].

Numerous mathematical models have been proposed to describe population movements in response to external navigation cues [25, 26, 27, 28]. Theoretical models of communication-based collective navigation are often individual-based random walk models [29, 30], where the behaviour of each individual is explicitly defined; though continuum models have also been proposed [8]. A common strategy has been to abstract interactions between individuals into a generic set of attraction, repulsion and alignment interactions [31, 10, 32, 33, 34, 35]. Interactions occur up to a maximum interaction range, or a defined number of neighbours [11]. Codling *et al*. employ this approach and demonstrate that group-based navigation is more efficient than individual navigation [10], provided that the environment is not highly turbulent. However, a widespread assumption in previous models of collective navigation is that an individual reliably perceives the behaviour of certain other individuals within the population. While this may be reasonable if the population is tightly clustered, it is less clear that this assumption holds for more dispersed populations. For example, in the presence of marine noise, the quality of perceived information may decrease markedly with distance [7]. Alternatively, for visual cues, the asymmetry of visual fields can restrict the directions from which a cue may be obtained, leading the spatial or topological structure of the population to impact on cue perception [36, 37, 38]. Whether a similar effect occurs for an auditory cue is less clear due to the physics of sound transmission through water. Another common assumption is that alignment interactions only rely on the (circular) mean of all observed headings to determine an optimal heading. Such an approach neglects all information regarding the variance in the observed headings. Consider a set of observed headings that are tightly clustered around the resultant mean heading compared to a set of observed headings, with the same resultant mean, but that are widely spread across all possible headings. It is plausible that the individual in the latter case would have less confidence in the resultant mean heading, compared to the first case. However, this reflection of decision making under uncertainty is typically not present in mathematical models. Certain approaches to modelling animal decision making between discrete options have been proposed previously [39, 40, 41]. In contrast, we seek to translate the uncertainty in the set of observed headings into uncertainty regarding a potential heading. It is currently unclear how the uncertainty in communication or perception may affect the ability of the population to undergo navigation due to the corresponding uncertainty in heading selection.

We develop a random walk model of communication-based collective navigation that incorporates uncertainty in the process of acquiring external guidance information. The random walk is biased according to a combination of observations of the heading of other individuals, balanced against the navigation information inherent to an individual. We examine the navigation performance of individuals governed by this model in a range of idealised information fields that represent the natural variability of navigation information. We demonstrate that communication results in a significant increase in navigation performance, provided that an individual can observe sufficiently many other individuals to overcome the uncertainty in communication. This increase in performance is most pronounced in the presence of information blindspots. To illustrate the utility of the framework we consider a case study of minke whale migration through the North East Atlantic Ocean, and examine how increased ambient noise due to drilling and other anthropogenic sources may inhibit migration.

## Methods

Consider an individual labelled *i* located in a two-dimensional plane at position **x**_*i*_(*t*) and navigating towards a target at **x**_target_. We model the individual’s movement path as a velocity jump random walk [42], an alternation between fixed velocity *runs* and *reorientations*. The duration of each run is sampled from an exponential distribution, parametrised by a turning rate parameter *µ*. For simplicity we impose quasi-instantaneous reorientation events, a constant turning rate and a fixed speed, *s*; these assumptions can be relaxed when tailored to a specific case study. For clarity of presentation, we presume a scaling such that *s* = 1. Velocities here are therefore unit vectors equivalent to an individual’s heading, and the formulae can be trivially modified for the case *s* ≠ 1. An individual with a heading represented by angle *θ*_*i*_, and corresponding velocity **v**_*i*_, moves according to

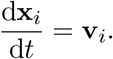

Navigation is encoded through the selection of a new heading at reorientation. We assume that this selection depends both on the *inherent information* available to an individual based on its current location, and on the *group information* obtained through communication with (or the perception of) other individuals (Figure 1). Model complexity is minimised by neglecting both repulsion and attraction, and we note that their effects have been considered in previous models [10]. We also assume that the post-reorientation heading is independent of the pre-reorientation heading. New headings are sampled from a von Mises distribution [43],

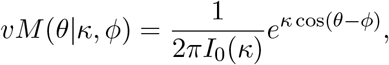

where *I*_0_(*κ*) denotes the modified Bessel function of order zero. The von Mises distribution is parametrised by a location parameter, *ϕ*, and a concentration parameter, *κ*. The location parameter reflects the most likely heading and an increasing concentration parameter increases the certainty of it being selected.

**Figure 1:**
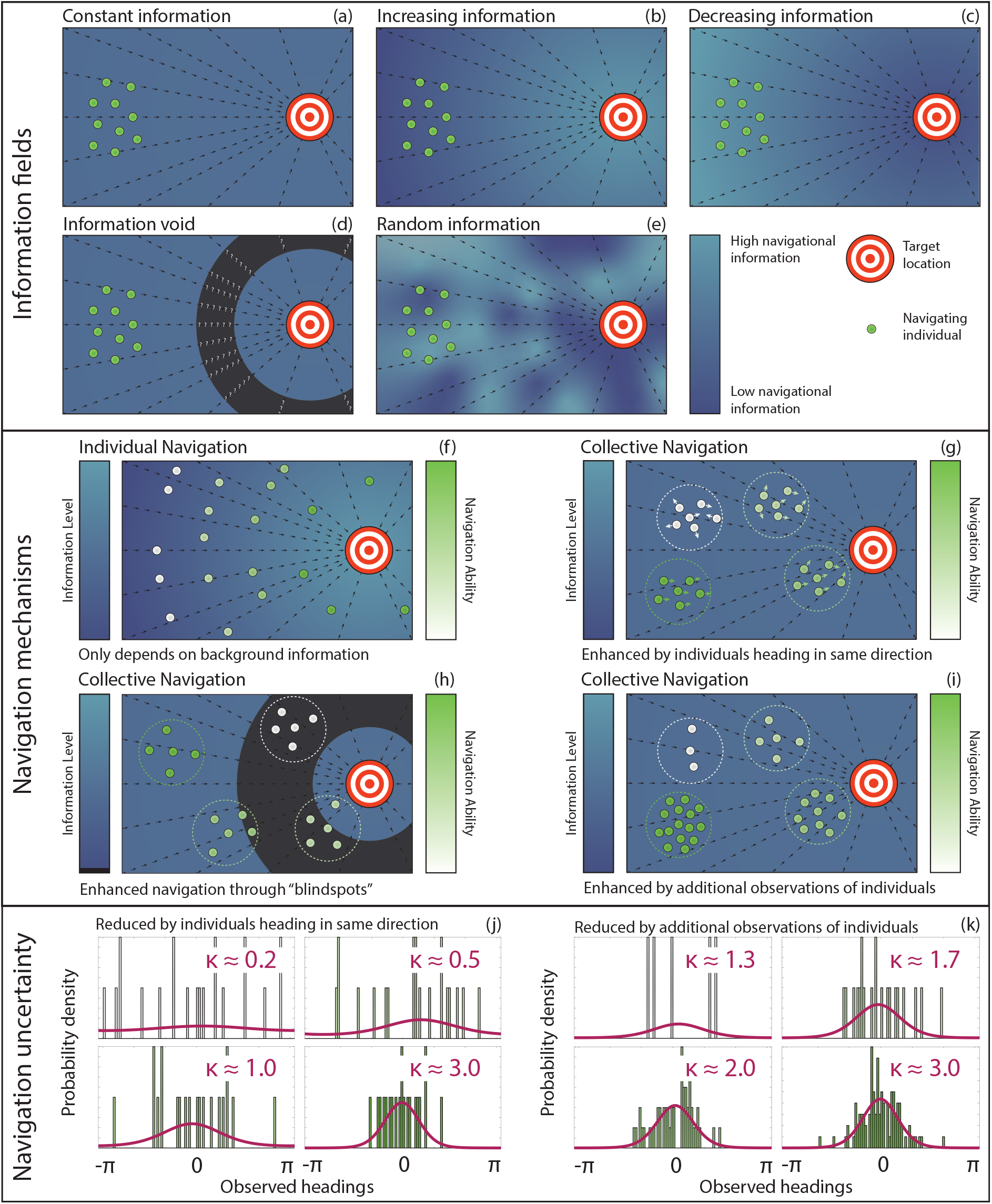
(a)-(e) Different types of information fields. The information field can either (a) be constant, (b) increase or (c) decrease as an individual approaches the target location, (d) contain a region of negligible information or (e) be randomly distributed. (f)-(i) Schematic highlighting the differences between (f) individual navigation and (g)-(i) collective navigation. For individual navigation, differences in navigation ability arise due to differences in the local inherent information. For collective navigation, differences in navigation ability may arise from (g) observing other individuals heading in similar directions, (h) observing other individuals in higher information regions, or (i) observing more individuals. We illustrate how this increase in navigation performance occurs by presenting (j),(k) the von Mises distributions (magenta) inferred from sets of observed headings that (j) are clustered to various degrees around a central heading or (k) include different numbers of observations. Distributions that are concentrated around the peak indicate decreased navigation uncertainty, as an individual is more likely to move in the direction of the target. Colours in the histograms indicate navigation ability, as in (f)-(i).

### Inherent information

Inherent information refers to the knowledge obtained when an individual samples navigation cues at its current position. The type of cue, an organism’s sensory processes, and the environment could all impact on the strength of this information. We assume inherent information is incorporated according to the von Mises distribution, where the concentration parameter *κ* is given by Ω(**x**_*i*_(*t*)). Therefore, Ω(**x**_*i*_(*t*)) defines the strength of the inherent information field for an individual currently located at **x**_*i*_(*t*). We assume that the location parameter is given by arg(**x**_target_ *-***x**_*i*_(*t*)); that is, the distribution resulting from inherent information is centred around the direction of the target location. We consider a range of inherent information fields, as presented in Figure 1, which are defined mathematically as

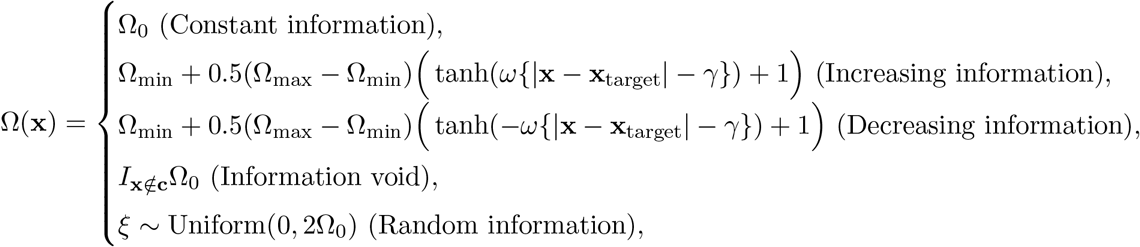

where Ω_0_ is the background information value for the constant information field. For the increasing and decreasing information fields, Ω_min_ and Ω_max_ are the minimum and maximum information values, respectively; *γ* is the distance from the target of the midpoint information value, and; *ω* is the information slope parameter. For the void information field, **c** is the set of locations inside an information void, and; *I* is an indicator function, equal to one if an individual is located outside of the information void, and zero otherwise.

### Group information

Under collective navigation, the individual’s inherent information can be enhanced by other individuals attempting to travel toward the same target. This information transfer can occur nonlocally through mechanisms that involve auditory, visual or other forms of communication. Consider an individual that perceives *n* other individuals with velocities **v**_*j*_, *j ∈* [1, *n*]. Each observation can be regarded as a sample from a von Mises distribution. This distribution will have similar parameters to the von Mises distribution sampled by individual *i*, presuming the maximum communication distance is not large compared to the length over which the background information changes. As such, we can construct the maximum likelihood estimates (MLEs) of *ϕ* and *κ* for the von Mises distribution governing the heading of the observed individuals. Of course, we are not suggesting that the individual is actually calculating the MLEs; rather, it is a convenient way of converting observations into a measure of average behaviour and certainty regarding that average behaviour. The MLE for the location parameter 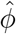 is simply the argument of the sum of the observations [44]:

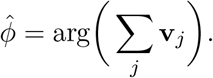

As individuals select a heading that is informed by the headings of other individuals this represents an alignment-type interaction, where individuals each seek to travel in the same direction.

Other possible interactions include attraction or repulsion [10, 45], where the individual moves towards or away from the centre of the observed neighbours. Attraction-type interactions may be particularly relevant in the final stages of migration, where an individual could signal its navigational success to the remainder of the population upon reaching the destination. However, as alignment-type interactions are relevant throughout the long migration process, we restrict our focus to these interactions at present. Further, including additional model complexity runs the risk of confounding or obscuring the relationship between uncertainty in observed headings and navigation performance.

It is plausible that information regarding an observed individual’s heading becomes distorted due to the distance the information travels before reaching the decision-making individual. This can happen, as an example, due to ambient noise disturbing an auditory signal. To account for this, a weighting kernel, *K*(*r*), can be used to describe the relative weighting placed on a signal as a function of distance between individuals, and hence the MLE for the location parameter will be

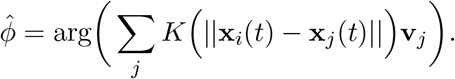

The MLE for the concentration parameter, 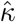, requires the following inverse problem to be solved, either via an approximation or through numerical techniques [44],

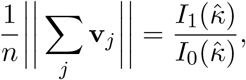

Where 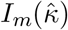 denotes the modified Bessel function of order *m*. If all observations are in a similar direction, then the MLE of the concentration parameter will be large, which implies that the individual has a high level of confidence in the location parameter. We can similarly include a weighting kernel to describe the relative weighting of information, which results in the MLE for the concentration parameter arising from

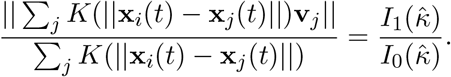

Here we will restrict our choice of weighting kernel to the Heaviside function

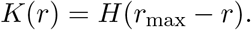

This implies that an individual places equal weight on observations of all individuals within a radius, *r*_max_, but ignores all other individuals. We refer to this radius as the *perceptual range*, the maximum distance over which a signal or cue can be perceived, for example mimicking the maximum perception distance relevant to communication through auditory and visual signals. Other natural choices for *K*(*r*) could include an exponential or power-law decay with distance, see for example [46]. Perceptual error could be directly imposed, as per previous studies [10], via the addition of a noise term to each observed heading. The strength of this noise term may be constant, or depend on a variable such as distance. Here we focus on uncertainty that arises organically due to the discrepancies present in the set of observed headings.

### Combining inherent and group information

The question remains regarding how to combine observed headings acquired nonlocally from the group with the inherent information available to an individual. Here we simply assume that the individual weights the observed headings against its inherent information:

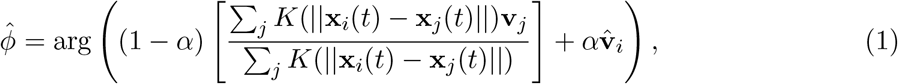

where *α* is the relative strength that an individual places on its inherent information with respect to heading and 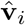 is the heading sampled from the distribution corresponding to the individual’s inherent information. If *α* = 0, the individual neglects inherent information and follows the crowd. If *α* = 1, the individual relies solely on inherent information. If *α* = 1*/*2, the individual places equal weight on inherent and group information. If *α* = 1*/*(*n* + 1) then the individual places equal weight on each observed individual, including itself. Another possibility is to allow *α* to vary according to, for example, the social relationships between individuals [17], such as for knowledgeable whale matriarchs, or the number of observed neighbours. However, for simplicity, here we restrict ourselves to constant *α* values for the entire population.

A similar weighting approach can be taken with the MLE estimate of the concentration parameter:

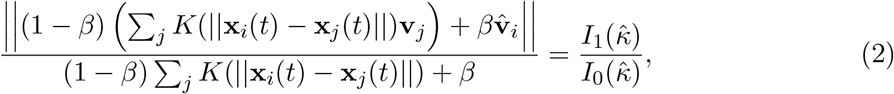

where *β* is the relative strength that an individual places on its inherent information with respect to concentration. The parameters *α* and *β* can be considered as the relative strength of social connections between the individual and its observed neighbours [17]. An investigation into the influence of the choice of *α* and *β* indicates that an approximately equal weighting between inherent and group information results in optimal navigation performance across a broad spectrum of information fields (Supplementary Information).

### Concentration parameter estimation

As detailed in Equation (2), the MLE for the concentration parameter can be calculated from *n* observations obtained from a von Mises distribution [43]. In [47] it is noted that this estimate is biased for either small *κ* or *n* values, both of which are likely to occur in group navigation. The authors proposed a correction which provides a less-biased mean, but exploring its distribution (compared to the uncorrected estimate) reveals that the reduction in bias is partly achieved through mapping the MLE of a large number of samples to zero (Supplementary Figure 1). While reducing the bias of the MLE for the concentration parameter is important, this reduction therefore occurs at the expense of a severely distorted distribution.

We therefore propose an alternative approach, where we repeatedly generate samples for fixed *n* and *κ* (i.e. a set of *n* headings obtained from the von Mises distribution with a concentration parameter *κ*) and determine the (uncorrected) 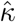 value for each sample. We repeat this process for a wide range of *n* and *κ* values and construct the distribution of *κ* values that give rise to a specific 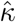 value for a fixed *n* value, which can be considered as the likelihood function, 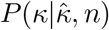 We pre-calculate a look-up table of 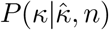 for 1 *≤ n ≤* 25 and 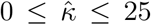, which addresses the issues associated with both insufficient observations and small *κ* values. In the model, an individual is informed by *n* observations and calculates 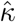 via Equation (2). We then sample from the likelihood function 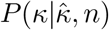 to provide an estimate of the concentration parameter of the von Mises distribution of the observed data. A comparison with the distributions from the uncorrected and corrected maximum likelihood estimate reveals that the likelihood function approach reduces the bias present without significantly inflating the number of estimates of the concentration parameter that are zero (Supplementary Figure 1).

### Model simulation

Initially, we distribute 100 individuals within a square of size 40, uniformly at random. The centre of this square is located at a distance of 300 from the target location. Initial headings are sampled according to the local inherent information. Individuals undergo motion at a fixed velocity until a reorientation event occurs. During a reorientation event, individuals undertake a three step process:

- First, an individual samples a heading from a von Mises distribution where the distribution parameters are informed by the inherent information;
- Second, an individual infers von Mises distribution parameters from a weighted combination of its sampled heading and the observed headings of the neighbours within its perceptual range via Equations (1) and (2), and;
- Third, the individual samples a heading from this inferred von Mises distribution and undergoes motion in the newly-sampled direction. Note that the second and third step only occur if there are other individuals within the perceptual range; otherwise the originally-sampled heading is retained.

When an individual arrives within a distance of 10 from the target location it is considered to have successfully navigated to the target, and is removed from the system. Unless stated otherwise we assume an implicit rescaling such that *s* = *µ* = 1. Consequently, the minimum *mean migration time* is *∼* 280 time units, which occurs if all individuals move in a straight line towards the target. Therefore, over the course of this journey each individual will, on average, re-evaluate its environment for navigation information several hundred times. For each simulation we track the number of individuals yet to reach the target location, the average number of neighbours within an individual’s perceptual range, and the average distance to the target of the individuals yet to reach the target location. All simulations are performed in Matlab R2020b, with the CircStat Toolbox employed for the necessary circular statistics [48].

## Results and Discussion

We first investigate navigation in a constant information field for a suite of perceptual ranges, and present the results in Figure 2. A perceptual range of zero corresponds to an individual navigating via inherent information only. Notably, we do not observe a monotonically increasing relationship between perceptual range and navigation ability, similar to the results observed by Codling *et al*. [10]. Rather, small perceptual ranges reduce navigation performance (compared to purely local navigation) and an improvement only occurs above a certain threshold. Examining the average number of neighbours offers insight into the root of this phenomenon. Perceptual ranges of five yield fewer than five neighbours throughout the simulated migration. This implies that relying on relatively few observations reduces navigation ability due to the uncertainty present in that small set of observations. If we consider the heading selection mechanism in the model, navigation using only inherent information corresponds to a single sample from a von Mises distribution centred around the heading of the target site, for a specified concentration parameter. In contrast, when group information is incorporated, navigation corresponds to a sample from an inferred von Mises distribution, constructed from a weighted combination of the aforementioned target heading sample and the headings of observed neighbours. The inferred distribution is not necessarily centred around the heading of the target, and for few observations the increase in the concentration parameter above background is insufficient to compensate for the increased variance in the location parameter. The decrease in performance is ameliorated by the presence of additional individuals within the perceptual range, as extra observations provide both a more reliable estimate of the heading of the target location and a further increase in the concentration parameter, ultimately improving navigation performance.

**Figure 2:**
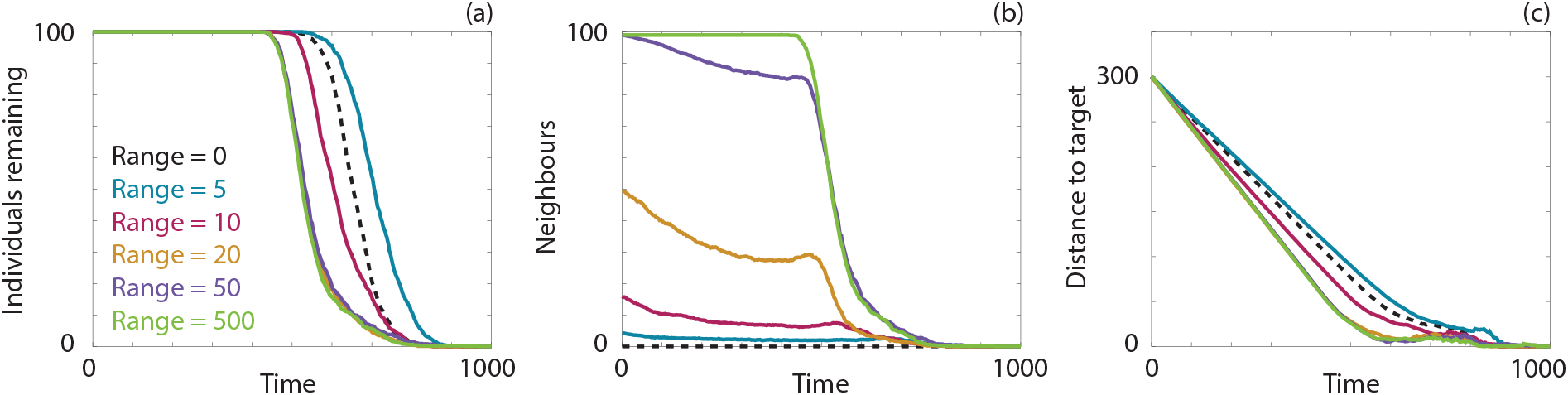
Navigation behaviour for 100 individuals in a constant information field. (a) The number of individuals remaining in the system over time. (b) The average number of neighbours within the perceptual range. (c) The distance between the target location and the average location of the individuals. Results are presented for a perceptual range of 0 (black, dashed), 5 (blue), 10 (magenta), 20 (orange), 50 (purple) and 500 (green). Parameters used are *α* = *β* = 0.5 and Ω_0_ = 1. All data are the average of ten realisations of the model.

We observe in Figure 2(a) that a perceptual range of 10, corresponding to between 10 and 20 neighbours, results in navigation performance that is more efficient than local navigation. The benefit of additional neighbours appears to plateau around 30 neighbours, as perceptual ranges of 20, 50 and 500 (effectively perceiving the entire population) demonstrate a similar navigation ability. We observe this phenomena across a suite of background information levels and perceptual ranges (Supplementary Figure 2). The nonmonotonic relationship between navigation performance and perceptual range has been observed previously [10]. However, in such models, where uncertainty is a predetermined parameter, the relationship becomes monotonic for sufficiently low uncertainty [10]. This is in contrast to our model, where uncertainty arises organically from the set of observed headings, and the nonmonotonic relationship is present for all investigated levels of background information.

As a further test we consider a set of simulations in which the number of individuals, *N*, and *r*_max_ are both altered such that 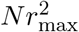 is kept constant. This ensures that approximately the same number of observed neighbours are within the perceptual range throughout each simulation. The resulting simulations (Supplementary Figure 4) show that the *proportion* of remaining individuals remains similar for each perceptual range considered, corroborating our hypothesis that the number of observed neighbours is the critical measure that informs navigation ability.

We next investigate navigation across the random, increasing and decreasing background information fields, illustrated in Figure 1. We calculate the time taken for 90% of the population to reach the target location for these fields, as well as for three constant information fields, and summarise the findings in Figure 3 (for detailed statistics, see Supplementary Figures 2 and 3). Unsurprisingly, increasing the background level of information improves navigation performance. Navigation behaviour for the random information field is similar to that observed for the constant information field with the same mean information level, suggesting that local fluctuations in background information do not significantly impact navigation ability compared to the mean background information. The expected nonmonotonic relationship between perceptual range and navigation ability is present in all cases. This relationship is less distinct for the decreasing information field and the higher constant information field, as the higher background information allows the population to remain together. In contrast, there is a stronger relationship between the perceptual range and the navigation ability for the increasing information field and the lower constant information field, due to the reduced background information, and corresponding population dispersal.

**Figure 3:**
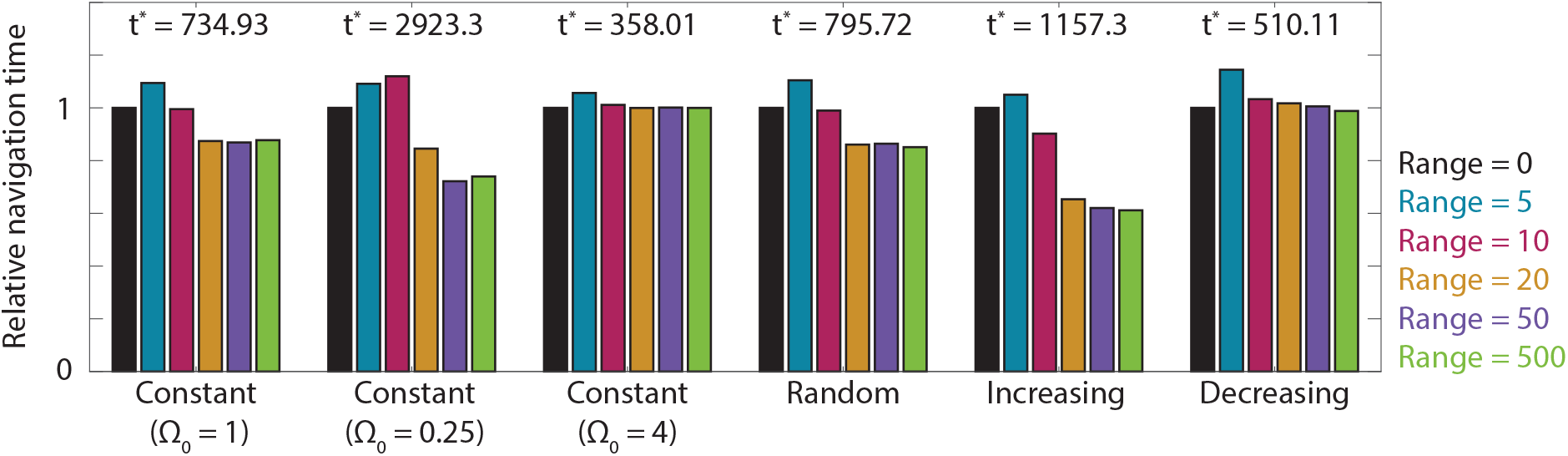
Navigation behaviour for 100 individuals in three constant information fields, a random information field, an increasing information field and a decreasing information field. Relative navigation time is defined as the time taken for 90% of the population to arrive at the target location, compared to navigation based on purely inherent information (denoted here as *t*^*∗*^). Results are presented for a perceptual range of 0 (black), 5 (blue), 10 (magenta), 20 (orange), 50 (purple) and 500 (green). Parameters used are *α* = *β* = 0.5 and Ω_0_ = 1, Ω_0_ = 0.25, Ω_0_ = 4, Ω_0_ = 1, and Ω_max_ = 2, Ω_min_ = 0.5, *ω* = 1*/*50, *γ* = 50, respectively. All data are the average of ten realisations of the model.

### Void information fields

We now turn our attention to void information fields, which describe a migration route that involves passage through one or more regions with negligible navigation information. We consider two forms of void information field: *chasm* fields, which contain a single void of fixed width that must be traversed en-route to the target, and; *patchy* fields, which contain multiple voids of variable size and shape. We first consider the chasm information field, and present the navigation behaviour in Figure 4. Notably, navigating using inherent information only becomes ineffective. Considering the distance between the centre of the population and the target location (Figure 4(d)) we observe that navigation is effective until reaching the region of zero information. At this point, unbiased random motion is required to navigate through the zero information region. Upon reaching the target-side of the void, an individual once more receives non-negligible information concerning the location heading.

**Figure 4:**
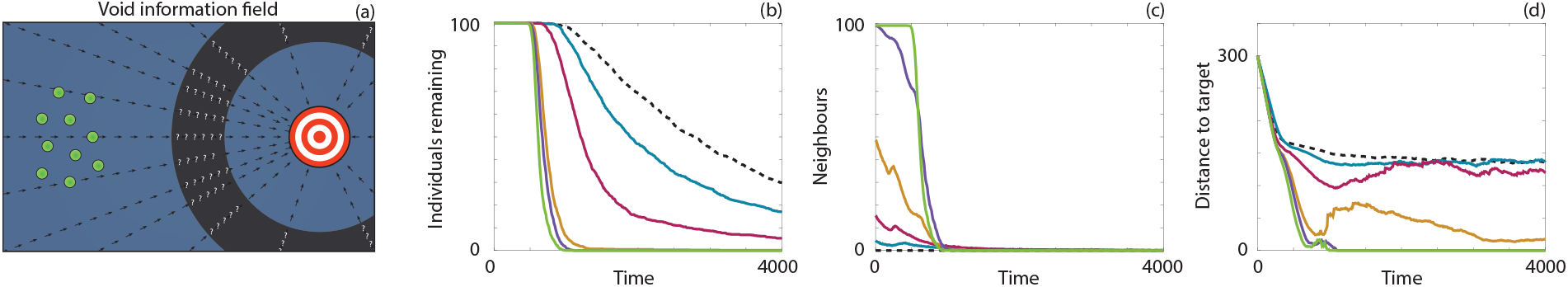
Navigation behaviour for 100 individuals in a chasm information field. (a) A schematic representation of the field. (b) The number of individuals remaining in the system over time. (c) The average number of neighbours within the perceptual range. (d) The distance between the target location and the average location of the individuals. Results are presented for a perceptual range of 0 (black, dashed), 5 (blue), 10 (magenta), 20 (orange), 50 (purple) and 500 (green). We note that the levelling off of the distance to the target statistics under low perceptual ranges is due to a few individuals remaining trapped in the information void. Parameters used are Ω_0_ = 1, **c** = {**x** | 125 *≤* ||**x** *-***x**_target_|| *≤* 175}, imposing a void of fixed width 50, and *α* = *β* = 0.5. All data are the average of ten realisations of the model.

In contrast, individuals that incorporate group information can observe individuals that may be outside of the zero information region, conferring a non-zero level of information regarding the heading of the target location. Intuitively, increasing the perceptual range increases the navigation ability of individuals. For a void of width fifty, navigation ability does not dramatically improve for perceptual ranges larger than twenty, corresponding to a range where an individual almost always observes neighbours outside the void. Note that even where an individual is itself unable to observe a neighbour located outside of the zero information region, the neighbours that it does observe could themselves be observing such individuals. Thus, improved navigation follows via a relay of information from individuals outside to individuals deep inside the void.

To shed further light on this phenomenon we calculate the effective concentration parameter and the number of neighbours as a function of distance from the target location for three different perceptual ranges and three sizes of information void, and present the results in Figure 5. The effective concentration parameter at each distance to the target is obtained from fitting a von Mises distribution to the differences between individual headings (at that distance to the target) and the target heading. As expected, the effective concentration parameter decreases inside the void, and this decrease is ameliorated by larger perceptual ranges. For larger voids, a prolonged decrease in the effective concentration parameter is observed. Notably, this decrease is asymmetric around the void midpoint. This asymmetry appears to arise through a short-lived persistence in navigation performance in the random walk as the individuals initially move into the void. Eventually, nearly all individuals find themselves within the void and group navigation provides a reduced but, crucially, non-negligible benefit. Subsequently, individual motion approaches an unbiased random walk and the group becomes disperse. As some individuals emerge on the target-side of the void, information begins to flow through to the individuals still within the void, causing the effective concentration to increase as individuals approach the target-side of the void. Larger perceptual ranges cause this increase to occur earlier, again demonstrating the benefit of an increased perceptual range with respect to navigation performance.

**Figure 5:**
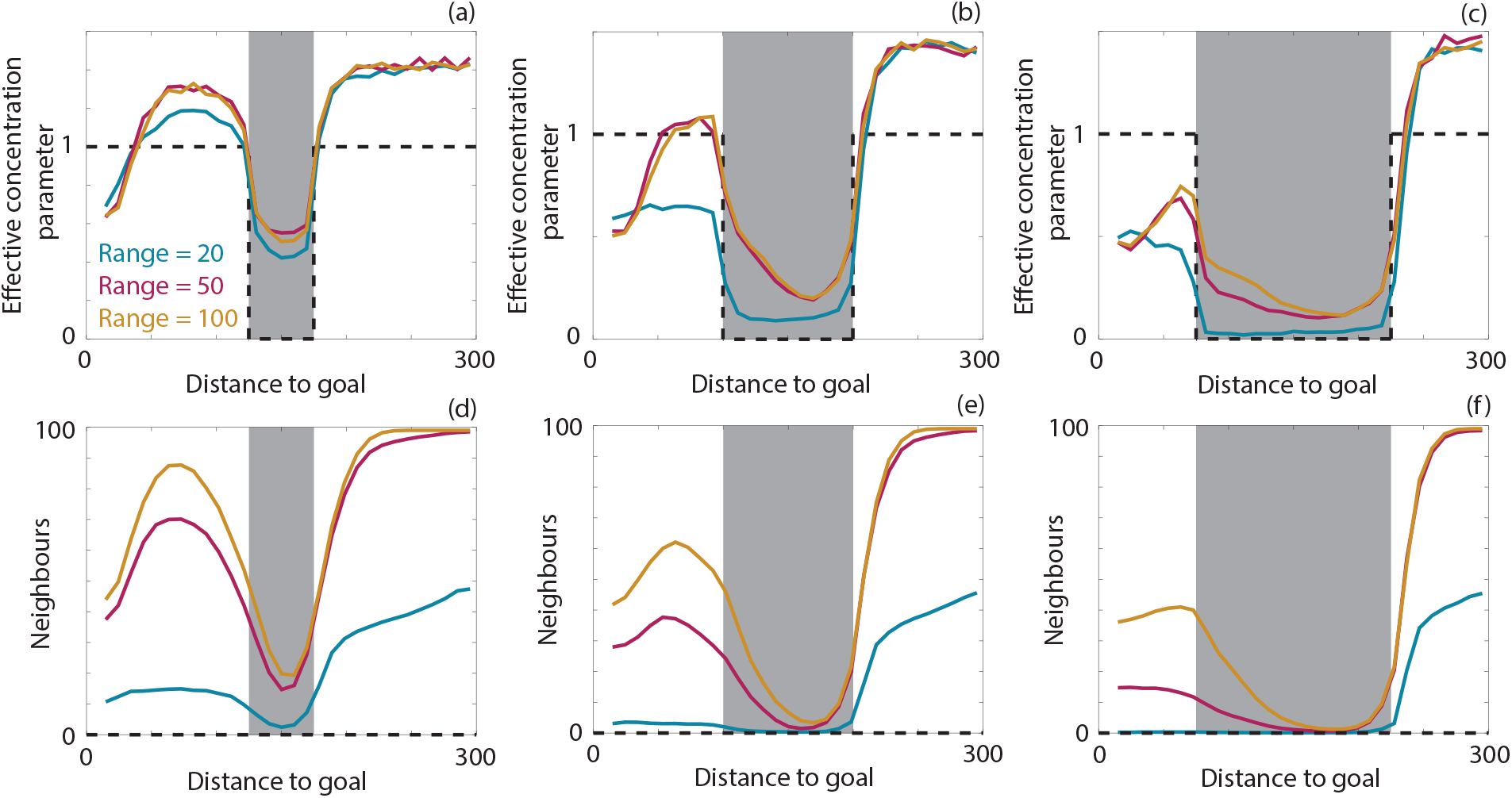
Effective concentration parameter and the average number of neighbours as a function of the distance to the target location for 100 individuals in a chasm information field with a void of width (a),(d) 50, (b),(e) 100 and (c),(f) 150 for perceptual ranges of 20 (blue), 50 (magenta) and 500 (orange). The results for purely local navigation correspond to the dashed black line. Parameters used are *α* = *β* = 0.5, Ω_0_ = 1 and (a),(d) **c** = {**x** | 125 *≤* ||**x** *-***x**_target_|| *≤* 175}, (b),(e) **c** = {**x** | 100 *≤* ||**x** *-***x**_target_|| *≤* 200}, (c),(f) **c** = {**x** | 75 *≤* ||**x** *-***x**_target_|| *≤* 225}. All results are the average of ten realisations of the model.

We next consider patchy void information fields. For the random information field in Figure 3, we treat randomness as a uniform random variable, sampled each time an individual undergoes reorientation. This can be interpreted as randomness at a fine scale, specifically at the length scale of the run between reorientations. It is equally plausible, though, to consider randomly-generated information fields that exhibit local correlation between information levels. That is, if an individual is in a low information area due to external factors, such as noise pollution, it is likely that the surrounding area is also a low information area. We generate such random information fields using a modified form of fractional Brownian noise; details are given in the Supplementary Information. Three representative fields are presented in Figure 6, where each field is generated following the same procedure but differences arise due to the number of nodes in the grid used to generate the noise. An increase in the number of nodes corresponds to finer structure present in the information field. For example, the information field in Figure 6(a) uses 16 times as many nodes in each direction as in Figure 6(i), and exhibits much finer spatial structure.

**Figure 6:**
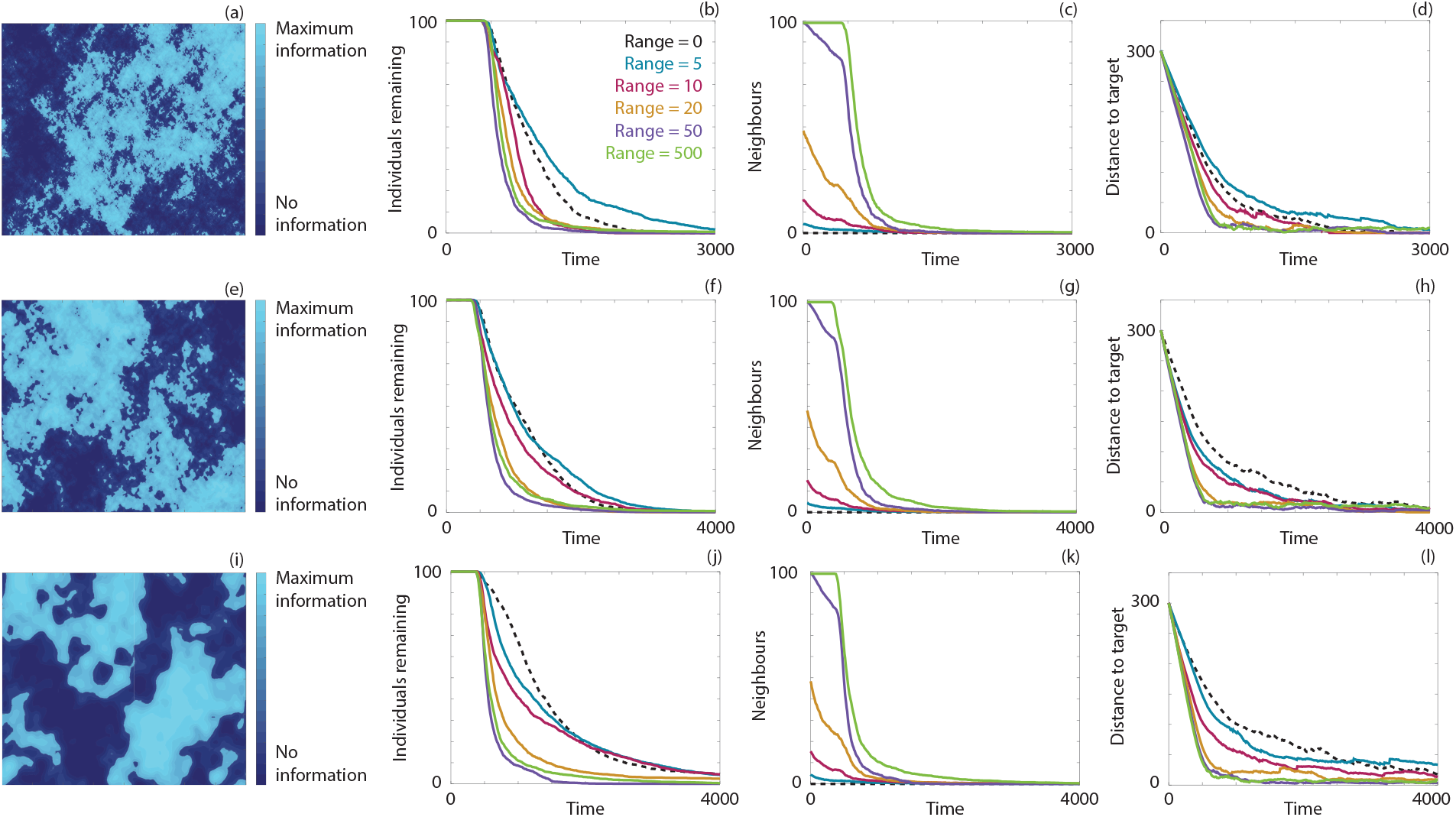
Navigation behaviour for 100 individuals in a fractional Brownian information field for different sizes of noise structures generated with (a)-(d) *m* = *n* = 400, (e)-(h) *m* = *n* = 100 and (i)-(l) *m* = *n* = 25 spatial nodes (see Supplementary Information). (a),(e),(j) Representative images of each randomly-generated field. (b),(f),(j) The number of individuals remaining in the system over time. (c),(g),(k) The average number of neighbours within the perceptual range. (d),(h),(l) The distance between the target location and the average location of the individuals. Results are presented for a perceptual range of 0 (black, dashed), 5 (blue), 10 (magenta), 20 (orange), 50 (purple) and 500 (green). Parameters used are Ω_0_ = 1 and *α* = *β* = 0.5. All data are the average of twenty realisations of the model.

For the three patchy information fields, with different scales of spatial structure, we calculate the navigation behaviour and present the results in Figure 6. Under the finest scale of spatial structure, which is closest to the original random information field, we again see the trend of increased navigation ability above a threshold number of observed neighbours. For coarser spatial structure, however, local navigation becomes less effective than the smallest perceptual range. For such information fields individuals can become trapped in low information areas, and therefore rely on random motion to return to high information areas. The risk of becoming trapped decreases with an increased perceptual range, as an individual can observe the movement of neighbours in high information areas. For realistic environments, where regions of low information may be present due to a range of external factors, this highlights the benefit of employing group navigation.

### Case study

To illustrate our approach, we consider migration through real world environments subject to different levels of anthropogenic noise. Specifically, we consider movements of minke whales (*Balaenoptera acutorostrata*) through portions of the North East Atlantic Ocean, surrounding the British Isles. Minke whales are the most frequently observed whale in these waters, found west of Ireland, off the north and east of Scotland and up to Iceland, Norway and beyond [49, 50, 51]. Sightings become less frequent in the southern North Sea, although seasonal aggregations have been observed in the Dogger Bank area near Denmark [52]. Notably, minke whales sightings remain largely confined to the April to October period and it is assumed that the population migrates south to winter in the mid Atlantic [53, 49]. We will consider two case studies of minke whale migration: first, a south to north migration through the North Sea from feeding grounds; second, migration through the East Atlantic Ocean, from southwest of Ireland to the west of Norway.

While minke whales are typically seen singly, or in pods of two or three, their vocalisations have been estimated to permit communication with conspecifics more than 100 km away [23]. Yet this calculation assumes a relatively “pristine” ocean soundscape, while modern marine environments are subject to significant anthropogenic activities that act to amplify ambient noise levels, such as shipping, wind farms, and oil exploration and drilling [20, 21]. Minke whale behaviour is strongly altered by ocean noise, for example through source avoidance [54, 55, 56] or raising call intensity [23]. The latter “Lombard effect” partially compensates for the noise, yet an *∼* 80% loss of their communication range has been estimated when ambient noise is raised 20 dB [23].

Motivated by the above, we construct an approximation of the noise levels present in the North Sea, in particular by exploiting the availability of offshore well location data (data obtained from the UK Oil and Gas Authority [57]). Specifically, the soundscape is formed through a sum of Gaussian noise profiles centred at each site. This is of course a simplification of the noise levels in the North Sea, as each offshore well may or may not be currently in operation, and we also do not include further significant noise sources, such as those due to shipping [58]. Consequently, this case study is primarily for illustrative purposes. Individual behaviour is modelled as detailed previously. However, to account for the presence of coastlines, jumps in the random walk that would result in an individual crossing onto land are aborted. Coastlines are constructed according to the Global Self-consistent, Hierarchical, High-resolution, Geography Database [59]. Under an ambient noise level of 65 dB it is estimated that minke whales are able to communicate up to 114 km [23] and it has been observed that minke whales travel at a speed of approximately 8 km/h [60]. In the absence of data, we make the assumption that minke whales undergo reorientation, on average, every 30 minutes.

In the first case study, we consider the change in minke whale navigation from a purported feeding ground in the North Sea [52] to a target location in the East Atlantic Ocean either in the presence of the anthropogenic noise in the North Sea, or in a pristine soundscape. The pristine soundscape corresponds to a constant level of background information available to the individuals. The information field arising from the offshore activity, as well as the navigation results, are presented in Figure 7. We observe that the migration occurs approximately 15% slower due to the presence of the noise pollution. This may impose a significant cost on the whales, who must expend additional energy to successfully navigate toward their target, reducing the energy stored for annual migration and breeding. Further, the noise pollution results in the group structure becoming dispersed. Close-knit group structure can be beneficial in terms of defence from predation [61], foraging [62], and, as we have demonstrated, for efficient navigation.

**Figure 7:**
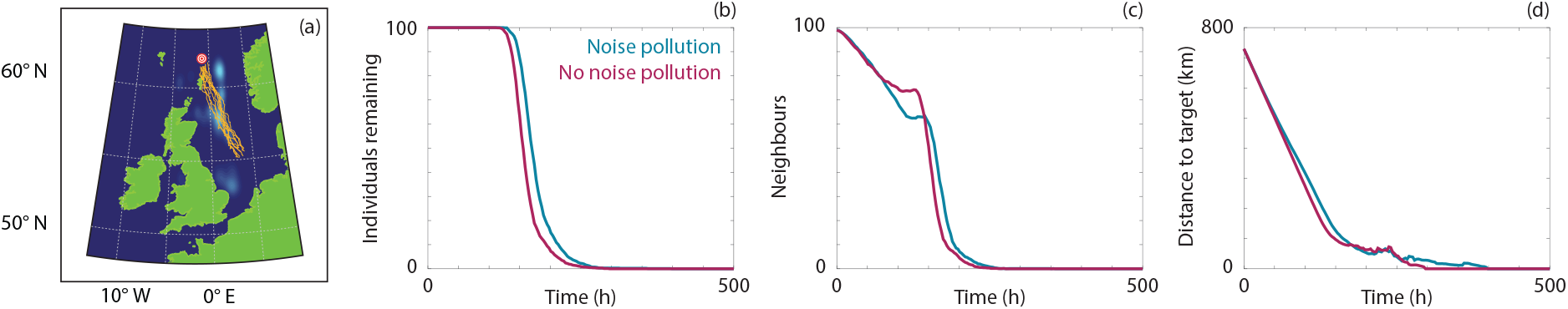
Comparison of migration from 55°N, 3°E to 62°N, 1°W in the North Sea for 100 individuals in the presence of noise from oil rigs (blue) or in the presence of pristine soundscapes (magenta). (b) The number of individuals remaining in the system over time. (c) The average number of neighbours within the perceptual range. (d) The distance between the target location and the average location of the individuals. Parameters used are Ω_0_ = 1, *α* = *β* = 0.5, *r*_max_ = 114 km, *s* = 8 km/h. Orange lines are a subset of simulated individual trajectories.

In the second case study, we consider potential increases in offshore noise pollution through the East Atlantic Ocean. We examine four different levels of noise pollution: the baseline case of approximately consistent noise; small-scale noise pollution, where 20% of the migration route contains significant noise pollution; medium-scale noise pollution, where 40% of the migration route contains significant noise pollution, and; large-scale noise pollution, where the entire migration route is enveloped by significant noise pollution. Sample trajectories under each noise pollution condition, as well as the navigation behaviour, are presented in Figure 8. Again, we observe that an increase in the total amount of noise corresponds to a decrease in navigation performance. Interestingly, the sample trajectories indicate that the whales somewhat avoid the areas of noise pollution, despite the model not containing any specific noise source avoidance behaviour. This is likely due to the decrease in target-oriented motion in the areas of noise pollution, resulting in random walks that cause the whales to leave the noisy area. Once outside of the area of noise pollution, the inherent information available to the whales increases and effective navigation toward the target location can take place. This highlights the need for areas with pristine soundscapes, where it remains possible to communicate and acquire inherent information effectively.

**Figure 8:**
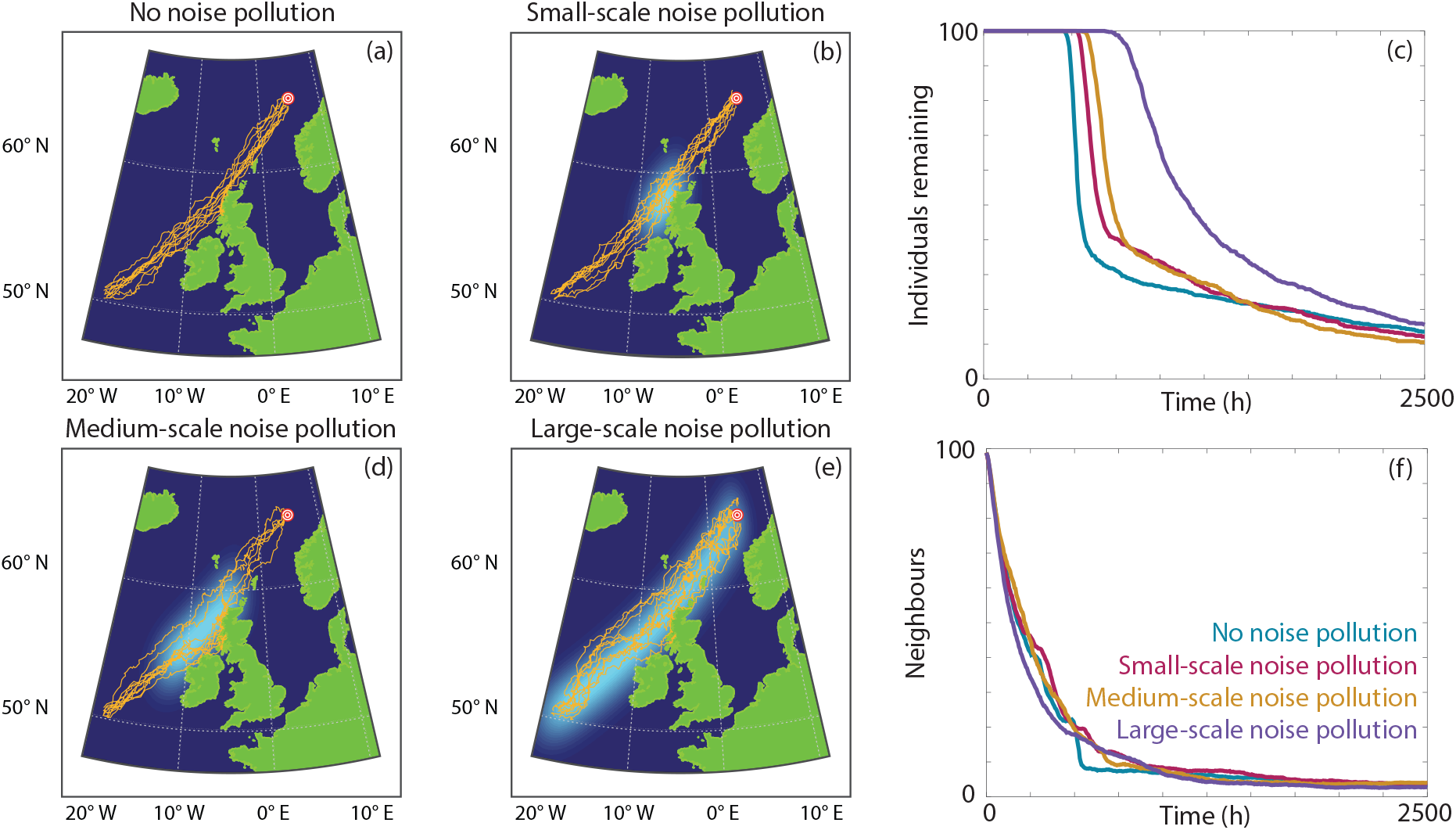
Comparison of migration from 50°N, 19°W to 65°N, 5°E in the East Atlantic Ocean for 100 individuals in the presence of (a) no noise pollution, (b) small-scale noise pollution, (d) medium-scale noise pollution or (e) large-scale noise pollution. (c) The number of individuals remaining in the system over time. (f) The average number of neighbours within the perceptual range. Parameters used are Ω_0_ = 1, *α* = *β* = 0.5, *r*_max_ = 114 km, *s* = 8 km/h. Orange lines are a subset of simulated individual trajectories.

## Conclusion

Migratory behaviour, conducted as a group across long distances, is a routine phenomena exhibited by many animal species [1]. The consistency of migration relies on the ability of animals to detect appropriate orienteering cues and/or to perceive group members that are migrating toward the same target. Both cue detection and perception can be inhibited by natural or anthropological phenomena, such as noise pollution. It is unclear how collective migration is impacted by this uncertainty in detection and communication. We have developed a novel mathematical model of collective migration and navigation that incorporates decision making under perceptual uncertainty. We employ this model to investigate how different information fields impact navigation performance, and illustrate the model via a case study application, specifically a disruption to minke whale migration due to anthropogenic noise pollution in the North East Atlantic Ocean.

We observe a nonmonotonic relationship between perceptual range and navigation performance for various information fields, in which increasing the maximum perceptual range from zero at first reduces, but subsequently improves, navigation performance, similar to previous studies [10]. Using group information raises certainty via increasing the concentration parameter, but at the expense of deviating the location parameter from the target heading. When the number of observed neighbours is low, the latter outweighs the former and navigation worsens. More observations tilts the balance the other way, and the intuitive improvement due to group navigation is observed. This is in contrast to previous studies, where the relationship between perceptual range and navigation performance is nonmonotonic above a threshold prespecified level of uncertainty, but monotonic below the threshold [10]. Our results suggest that the number of neighbours, rather than perceptual range, is the critical determinant as to whether group information improves navigation. Further, the critical perceptual range will depend on the degree to which the population remains clustered. Our model, though, has applied the “metric distance” approach to communication, i.e. permitting interactions with any number of individuals up to some fixed distance apart. This is a common assumption in models [11] and is consistent with, for example, visual or auditory systems, where there is an upper limit on the perceptual range. It is less certain, however, whether an individual can process more than a certain number of neighbours; the “topological distance” model postulates that an individual reacts to a fixed number of nearest neighbours, regardless of proximity [63]. This is more relevant for animals that are densely clustered, such as in starling flocks, where an individual could feasibly observe many other individuals within its perceptual range. Whether this alternative approach robustly (i.e. regardless of the form of navigational field and degree of clustering) generates improved navigation above a critical fixed interaction number remains to be explored.

Lengthy migrations towards a target can be divided into a set of stages [6]: a long distance phase, a homing phase, and a pinpointing phase. Often, distinct navigational cues will be used in the distinct phases [6]. For example, the mechanisms used by marine turtles to home on remote nesting beaches is believed to involve the geomagnetic field at longer distances, and olfactory or visual information at shorter ranges [64, 65, 66]. Within such a context, void information fields can be viewed as an information gap, where the navigator must cross some space of low information to bridge the regions where long and short distance cues are effective. Group navigation becomes particularly advantageous here, where for large-scale information voids any nonzero level of group information improves navigation. An observation acquired from a minimally more knowledgeable neighbour is sufficient to provide some target-oriented drift. Benefits of group navigation extend to very short perceptual ranges, that is, where the perceptual range is a full order of magnitude below the dimensions of the void region. Information reaches the centre of the void through a relay, so that an individual deep inside the void will still receive some information even when all of its observed neighbours are inside the void. This can occur as just one of those neighbours’ neighbours may be in a region of high inherent information.

Navigation can be disrupted by a decrease in inherent information (i.e. reduced quality of the external navigating cue) and/or a decrease in group information. As expected, the loss of either information source reduces navigation performance. Notably, though, a distinct response is observed according to the general level of inherent information: a loss of inherent information is more disruptive when the background navigation information is high, while a loss of group information has a more severe impact when the background navigation information is low (Supplementary Figure 8). This reinforces the notion that group navigation is particularly advantageous within weaker information environments and stems from the degree to which the population spreads: low (high) information environments leads to greater (lower) spreading and the average number of neighbours in the perceptual range is lower (higher). It is perhaps logical to suppose, therefore, that populations will have evolved different strategies for reducing the impact of different types of navigational disruption. This could occur, for example, by spreading out to the limits of their perceptual range when passing through low information regions and maintaining a tight/compact form when the perceptual range is inhibited. Against such a m strategy, spreading out could render the population vulnerable to unpredictable communication range loss, e.g. sudden noise sources. One plausible adaptive response to a noisy environment is to adjust the rate of heading selection. It has been observed that three-spined sticklebacks (*Gasterosteus aculeatus*) update their velocities more frequently in response to a perceived threat [45]. As such, one extension of our model is to relax the assumption of a constant turning rate, and to allow it to vary with, for example, noise levels. It is known that certain animals, such as night-migratory Eurasian blackcaps (*Sylvia atricapilla*), undertake random motion in the absence of an expected navigation cue [67]. However, another possible adaptive response to a noisy environment or an abrupt loss of navigation information is for the individual to continue on a similar heading [68]. That is, an individual makes smaller changes in heading due to an imposed correlation between the new heading and the previous heading [69, 68]. It would be instructive to compare migration paths for an individual that switches to a correlated random walk model when entering an information void, with those of our model, where an individual continues an unbiased, uncorrelated random walk; however, we leave this for future work. We note that our current model does not allow the population to control their separation through attraction/repulsion behaviour, and a natural extension is to adapt the model to include such behaviour [10]. Admitting control of group structure can facilitate investigations of whether particular group shapes are advantageous, such as an elongated shape to allow information transfer across voids. “Leader-type” individuals are also likely to be important for group structure, for example by adopting a specific spatial position with respect to the group to maximise information transfer. Such individuals could be incorporated by imposing larger *α* and *β* values for observed headings corresponding to “leader-type” individuals, signifying that such headings are more important than other headings.

We have restricted our attention to static information fields, that is, levels of inherent information that only vary in space. Temporal variation could occur due to, for example, weather conditions or intermittent human activity. The degree to which dynamic variation impacts on journeys will, naturally, depend on the duration of the disruption: the return of little penguins (*Eudyptula minor*) from daily foraging is delayed by heavy fog, possibly due to their reliance on visual navigating cues [70]. The impact of longer lasting dynamic variability could, to a degree, be inferred from the existing results. Slowly shifting cloud cover could be represented by evolving patchy void environments, and group navigation is always beneficial in such scenarios. Dynamic variability could also arise from the behaviour of individuals that have reached the target. The arbitrary modelling decision here has been to remove such individuals from the system, and hence those individuals no longer influence navigation. It is also possible that individuals actively communicate on arrival, providing information about the target location. Such behaviour has been suggested in humpback whale populations, where individuals sing upon reaching winter grounds, which may attract other humpbacks to the area [71]. This behaviour can be incorporated in our modelling framework by imposing an attraction-type interaction, rather than an alignment-type interaction, for individuals that have arrived at the target location.

Anthropogenic activity has substantially increased ocean noise levels over the past century [19, 20]. As a case study, we have explored noise-impacted minke whale migration through the North East Atlantic Ocean. Hypothesising that minke whales use vocal communication to share navigation information, we have shown that increasing noise pollution decreases navigation performance, thereby demanding longer travel times. Migration is costly, depleting an animal’s energy reserve without any guarantee of replenishment en-route, and hence any increased expenditure is disadvantageous to population fitness. Without an explicit representation of noise source avoidance, pathways are diverted from high noise areas as the animal searches for a route with adequate navigation information. Nevertheless, it remains important to stress that this study has been primarily expository in nature, and several further extensions demand consideration. First, as highlighted above, we have ignored the consequences of other behavioural interactions, for example an individual changing direction and speed to avoid a noise source, or regulating intergroup spacing. Second, we have ignored ocean currents, which could act to assist or hinder navigation and impact on sound propagation. Third, our incorporation of noise impacts has been rather simplistic: we neglect other potential sources, such as shipping, and we do not explicitly include the physics of noise propagation within the ocean. These caveats aside, the framework is highly adaptable, can be easily translated to other geographical locations, and can be extended in a modular fashion to include data inputs such as ocean currents, bathymetry and sound profiles.

## Data access

The code used to implement the mathematical framework is available at https://github.com/DrStuartJohnston/collective-navigation. The look-up table is available at https://melbourne.figshare.com/articles/dataset/kappaCDFLookupTablemat/14551614.

## Author contributions

STJ conceived the study, developed the modelling framework, designed and performed the numerical experiments, designed and performed the analysis, and wrote and edited the manuscript. KJP conceived the study, developed the modelling framework, designed the numerical experiments, designed and performed the analysis, and wrote and edited the manuscript. All authors gave final approval for publication.

## Supplementary Information

### Concentration parameter estimation

Comparing the likelihood function, 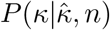, the distribution of the uncorrected maximum likelihood estimate, 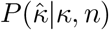, and the distribution of the corrected maximum likelihood es-timate, 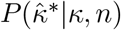, shows that both the corrected estimate and the likelihood function avoid the heavy tail present in the uncorrected estimate, Figure 9. However, the likelihood function approach reduces the bias present without significantly inflating the number of estimates of the concentration parameter that are zero.

**Figure 9:**
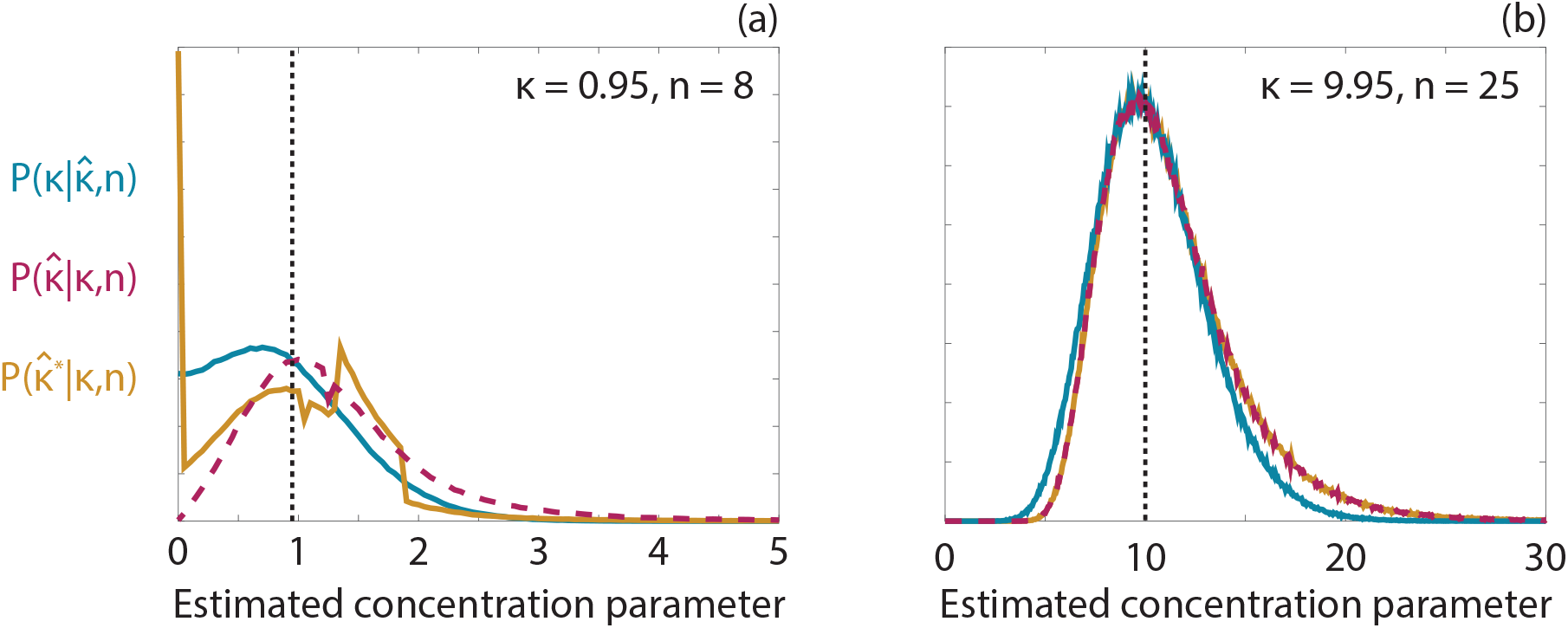
Comparison between the distributions for the *κ* likelihood (blue), 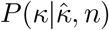, un-corrected MLE (magenta, dashed), 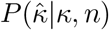, and the corrected MLE (orange), 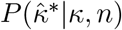, for (a) *κ* = 0.95 and *n* = 8 and (b) *κ* = 9.95 and *n* = 25. The black dashed line correspond to the true *κ* value. The corrected MLE proposed in Best & Fisher (1981) is given by 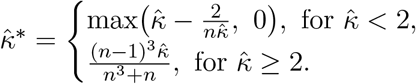.

### Different levels of background information

We repeat the analysis presented in Figure 2 (main manuscript) for a four-fold increase and a four-fold decrease in background information, and present the results in Figure 10. Broadly, we observe the same trend, where small perceptual ranges lead to both a reduction in the number of neighbours and greater uncertainty. For the lower level of background information, Figures 10(a)-(c), navigation performance remains degraded for even larger perceptual ranges. For example, compare the navigation performance for a perceptual range of 10 in Figure 10(b) to the corresponding perceptual range in Figure 2(b) (main manuscript). Lower background information results in an increase in the spread of the population, generating a precipitous drop in the average number of neighbours and, in turn, a decrease in navigation ability. For higher background information (Figures 10(d)-(f)) the population does not spread out to the same extent. As such, the number of neighbours within the perceptual range is higher than in both Figures 10(b) and 2(b) (main manuscript), and the change in navigation ability due to a small perceptual range is less pronounced. Interestingly, for the low level of background information (Figures 10(a)-(c)), a large number of neighbours within the perceptual range corresponds to a more pronounced increase in navigation ability (relative to purely local navigation), compared to higher levels of background information. This implies that in low information environments there is more benefit associated with maintaining a close-knit population structure, with respect to navigation, than in high information environments.

**Figure 10:**
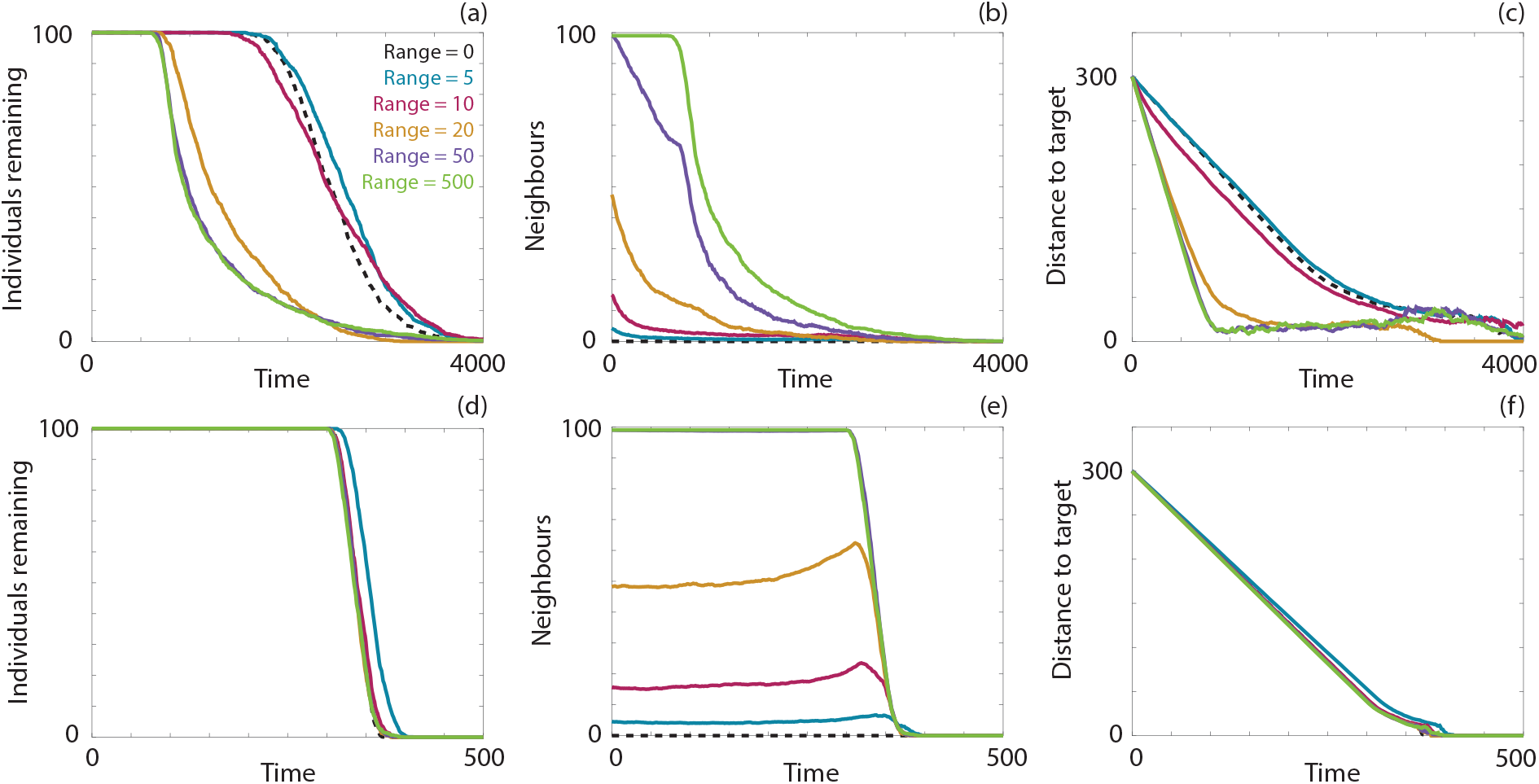
Navigation behaviour for 100 individuals in a constant information field. (a),(d) The number of individuals remaining in the system over time. (b),(e) The average number of neighbours within the perceptual range. (c),(f) The distance between the target location and the average location of the individuals. Results are presented for a perceptual range of 0 (black, dashed), 5 (blue), 10 (magenta), 20 (orange), 50 (purple) and 500 (green). Parameters used are *α* = *β* = 0.5 and (a)-(c) Ω_0_ = 0.25, (d)-(f) Ω_0_ = 4. All data are the average of ten realisations of the model.

### Varying information fields

We now consider navigation for the random, increasing and decreasing background information fields. In the random information field the inherent information is sampled from a uniform distribution on (0, 2Ω_0_). Navigation behaviour, presented in Figures 11 (a)-(d), is similar to that observed for the constant information field with the same mean information level, suggesting that local fluctuations in background information do not significantly impact navigation ability compared to the mean background information. Under increasing (Figures 11(e)-(h)) and decreasing (Figures 11(i)-(l)) information fields we again see that a small number of observed neighbours reduces navigation ability, compared to local navigation, and that observing sufficiently many neighbours enhances navigation ability. For the decreasing field the effect of perceptual range on navigation ability is not particularly strong, as the high information level at the beginning of the simulation allows the population to remain together. In contrast, when the increasing information field is considered, an initially low level of information leads to population dispersal. As such, there is a stronger relationship between the perceptual range and the navigation ability.

**Figure 11:**
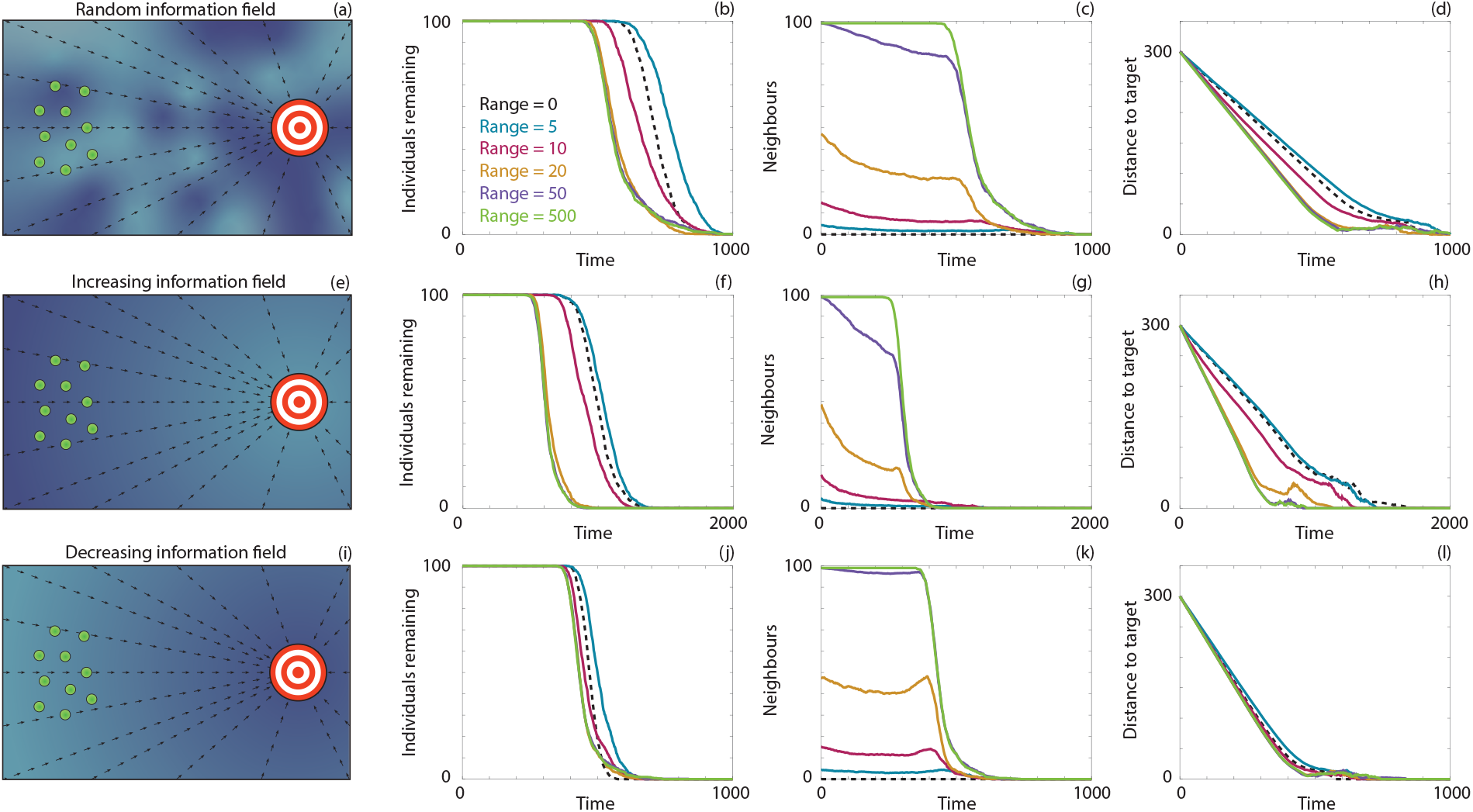
Navigation behaviour for 100 individuals in (a)-(d) a random information field, (e)-(h) an increasing information field, and (i)-(l) a decreasing information field. (a),(e),(i) A schematic representation of each type of field. (b),(f),(j) The number of individuals remaining in the system over time. (c),(g),(k) The average number of neighbours within the perceptual range. (d),(h),(l) The distance between the target location and the average location of the individuals. Results are presented for a perceptual range of 0 (black, dashed), 5 (blue), 10 (magenta), 20 (orange), 50 (purple) and 500 (green). Parameters used are (a)-(d) Ω_0_ = 1, (e)-(l) Ω_max_ = 2, Ω_min_ = 0.5, *ω* = 1*/*50, *γ* = 50 and *α* = *β* = 0.5. All data are the average of ten realisations of the model.

### Critical perceptual range

To clarify the relationship between the perceptual range, the number of observed neighbours and the navigation ability of the population, we present results where we vary the initial number of individuals, *N*, as well as the perceptual range, *r*_*max*_. Crucially, we select these values such that 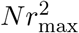 is constant. This is approximately equivalent to ensuring that there are the same number of observed neighbours within the perceptual range in each simulation. In Figure 12(a), we observe that for a perceptual range of ten, there are more individuals remaining in the simulation compared to larger perceptual ranges. However, if we consider the *proportion* of remaining individuals, presented in Figure 12(b), we see that the results are extremely similar for all three perceptual ranges. The number of observed neighbours is relatively consistent, as shown in Figure 12(c), as is the average distance between the target location and the population. These results corroborate our above hypothesis that the number of observed neighbours is the critical measure that informs navigation ability. However, it is not necessarily straightforward to determine the number of neighbours *a priori*, as it depends both on the background level of information, which influences population dispersal, and the perceptual range, which allows individuals to observe neighbours that are located farther away.

**Figure 12:**
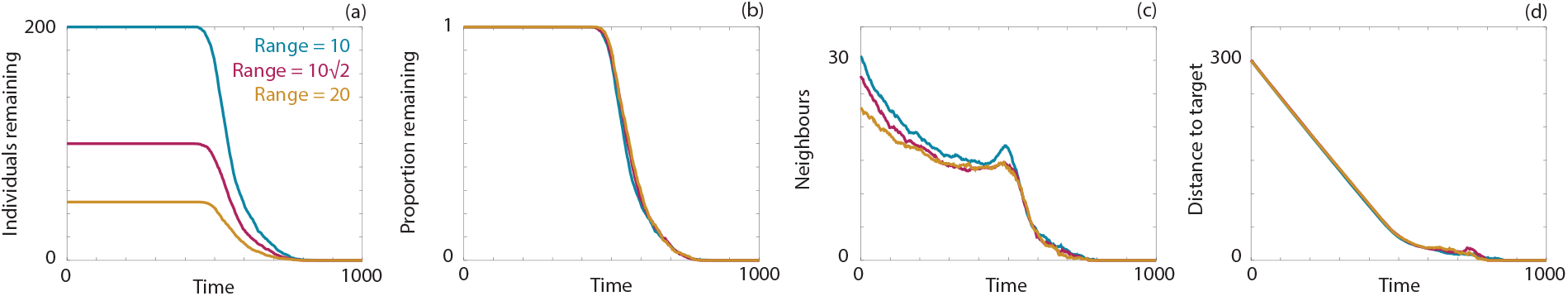
Navigation behaviour for 50 (orange), 100 (magenta) and 200 (blue) individuals in a constant information field. (a) The number of individuals remaining in the system over time. (b) The proportion of the initial number of individuals remaining in the system over time.(c) The average number of neighbours within the perceptual range. (d) The distance between the target location and the average location of the individuals. Results are presented for a perceptual range of 10 (blue), 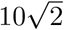 (magenta) and 20 (orange). Parameters used are Ω_0_ = 1 and *α* = *β* = 0.5. All data are the average of ten realisations of the model.

We extend our earlier analysis for a suite of perceptual range and background information combinations to determine the critical perceptual range, where collective navigation becomes more efficient than navigation based solely on inherent information. We calculate the time taken for 90% of the population to reach the target for a given perceptual range and background information level, relative to local navigation for that background level, and present the results in Figure 13. We observe a decreasing relationship between the critical perceptual range and the background information. This reinforces the previous observation that the benefit of a larger perceptual range is more pronounced for lower background information levels.

**Figure 13:**
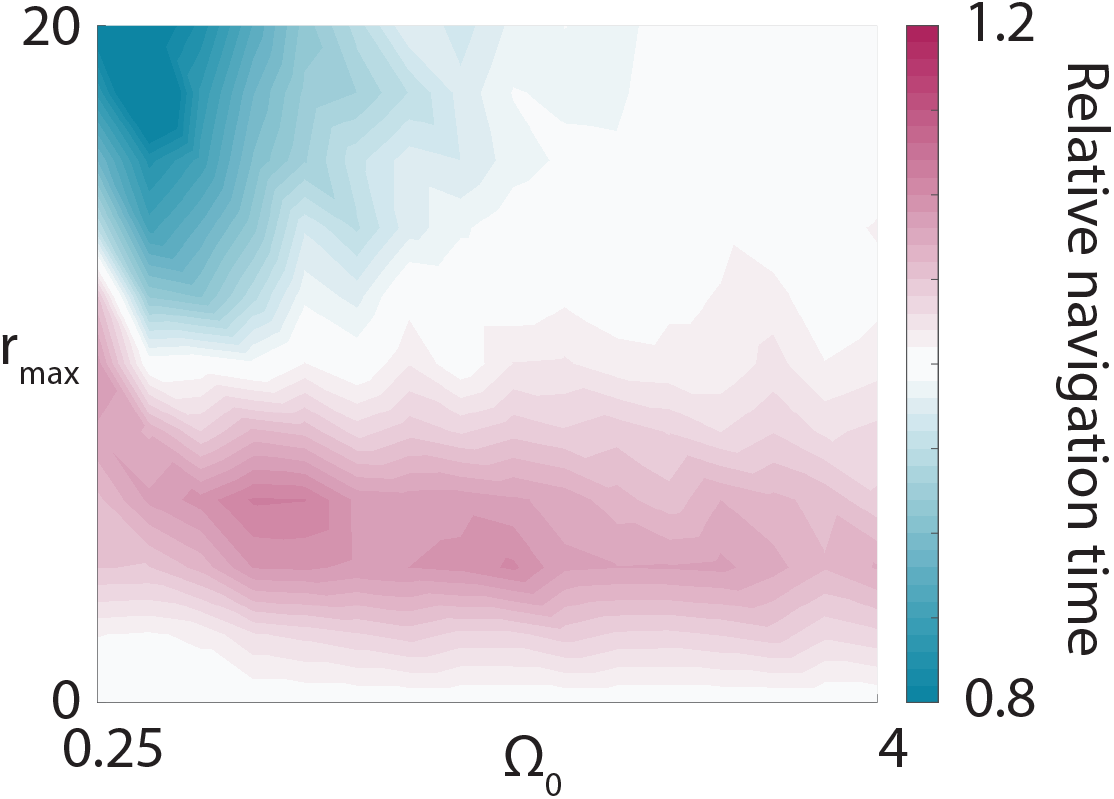
Relative navigation time required for 100 individuals in a constant information field for a suite of background information levels (Ω_0_) and maximum perceptual ranges (*r*_max_). Relative navigation time is defined as the average time taken for 90% of the population to reach the target location, compared to navigation based on purely inherent information, that is, *r*_max_ = 0. Here *α* = *β* = 0.5. All data are the average of ten realisations of the model.

### Balancing inherent and group information

Thus far we have assumed that individuals equally balance inherent and group information. It is also possible that individuals place different values on these information sources. We therefore consider a range of values for *α* and *β*, representing the weight placed on the inherent information for the heading and concentration parameters, respectively. For each parameter pair, we calculate the average time required for 90% of the population to arrive at the target location in a constant information field (Figures 14(a)-(b)). Neglecting inherent information, with respect to the concentration parameter (low *β* values), results in an increase in navigation time. This is due to the decrease in the inferred concentration parameter as less weight is placed on the inherent observation. Striking a balance between group and inherent information, with respect to the heading, reduces the navigation time. If an individual places too much weight on inherent information, it does not benefit from the averaging of headings that occurs when considering group information. Conversely, if an individual places too much weight on group information, it becomes difficult to undergo significant changes in heading, as each individual relies heavily on the directions of its neighbours. Figure 14(b) contains the subset of results in Figure 14(a) that satisfy *α* = *β*, that is, the same weighting is applied to both the heading and concentration parameters. We observe that the minimum navigation time occurs near the middle of the range. This is in contrast with the minimum navigation time in Figure 14(a), which occurs for (*α, β*) = (0.3, 1). However, this overall minimum arises due to the reduction in uncertainty when neglecting group information for the concentration parameter, as the distribution from which the concentration parameter is inferred from becomes singular around the heading as *β →* 1. As such, this may be an unrealistic assumption, as an individual would be equally confident about an inferred heading obtained from two observations, of directly opposite headings, as an inferred heading obtained from 100 observations clustered around a certain direction.

**Figure 14:**
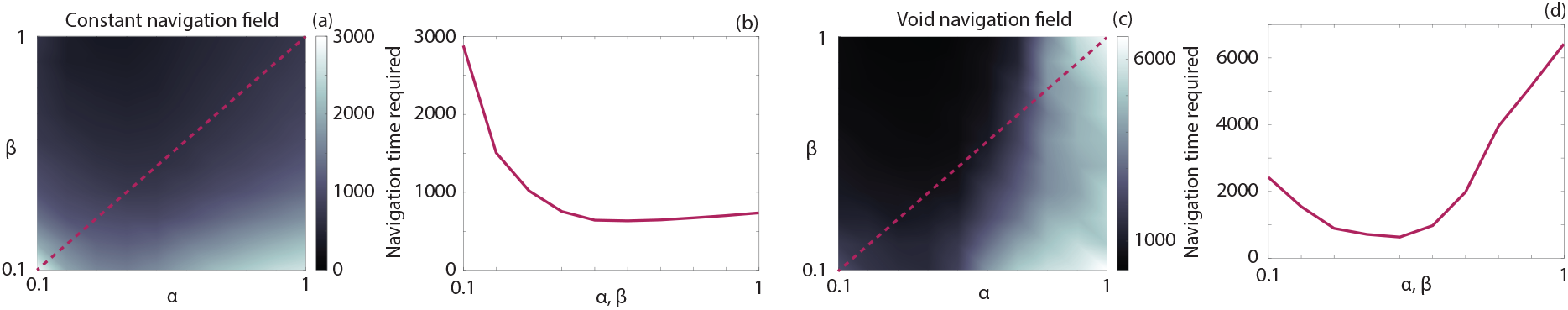
Navigation time required for 100 individuals in (a)-(b) a constant information field or (c)-(d) a void information field for (a),(c) a suite of (*α, β*) values, and (b),(d) a range of *α* = *β* values. Navigation time is defined as the average time taken for 90% of the population to reach the target location. Here Ω_0_ = 1, *r*_max_ = 20, and (c)-(d) **c** = {**x** | 125 *≤* ||**x** *-***x**_target_|| *≤* 175}. All data are the average of ten realisations of the model. The dashed line in (a),(c) corresponds to the results in (b),(d).

We perform a similar analysis for a chasm information field, and present the results in Figures 14(c)-(d). Compared to the constant information field, the region of poor navigation is now found to occur for large *α*. Within this regime, an individual places too much weight on its inherent information, with respect to the concentration parameter, which is clearly disadvantageous when the individual finds itself within a void. In Figure 14(d) we present the subset of results in Figure 14(c) that satisfy *α* = *β*. As for the constant field, the minimum clearly lies in the centre of the range. Further investigations for increasing or decreasing fields yield the same, suggesting an overall optimum strategy in which approximately equal weight is placed on inherent and group information.

We repeat the analysis in Figure 14 for both increasing and decreasing information fields and present the results in Figure 15. The observed navigation time across the *α* and *β* parameter space for both the increasing and decreasing information fields are qualitatively similar to the constant information field. Again, the minimum navigation time under the restriction that *α* = *β* arises when *α* = *β* = 0.5 or *α* = *β* = 0.6, suggesting that an approximately equal weighting between inherent and group information results in optimal navigation performance.

**Figure 15:**
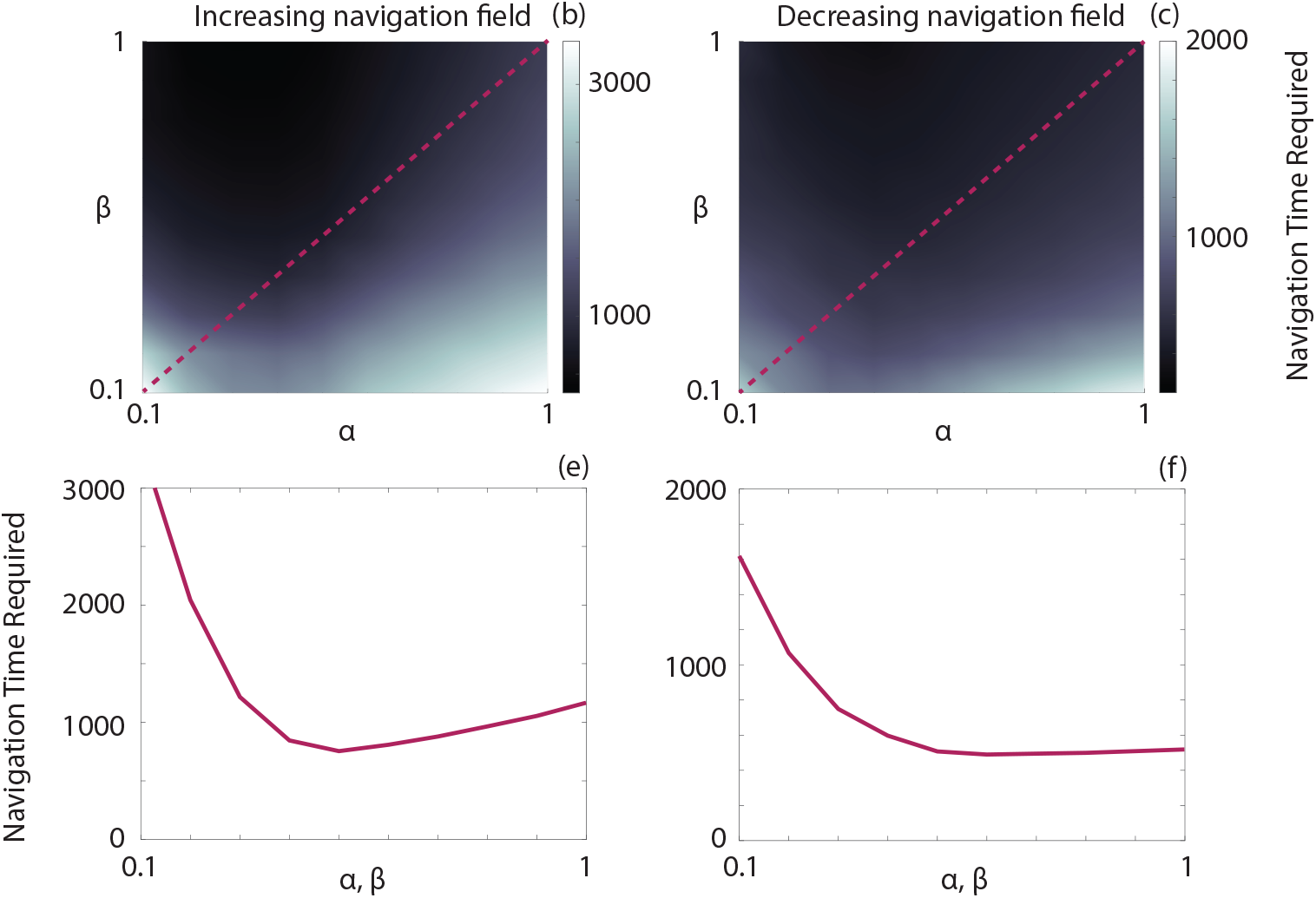
Navigation time required for 100 individuals in a (a),(d) void information field, (b),(e) increasing information field, and (c),(f) decreasing information field for (a)-(c) a suite of (*α, β*) values, and (d)-(f) a range of *α* = *β* values. Navigation time is defined as the average time taken for 90% of the population to reach the target location. Parameters used are (a),(d) Ω_0_ = 1, (b)-(c), (e)-(f) Ω_max_ = 2, Ω_min_ = 0.5, *ω* = 1*/*50, *γ* = 50. All data are the average of ten realisations of the model. All data are the average of ten realisations of the model. The dashed line in (a)-(c) corresponds to the results in (d)-(f).

### Range-reducing or information-reducing noise

For the void information fields above, disturbances are considered in the form of step-like changes to inherent information, so that an individual entering the void possesses negligible inherent information. It could also be appropriate to consider disturbances in the form of perceptual range reduction, i.e. an impaired individual can only perceive its neighbours up to a reduced distance. To test this we examine whether an individual can more efficiently undertake navigation where it cannot detect its neighbours due to noise, compared to where the individual loses the inherent information regarding the target location due to noise. Specifically, an individual entering a void region is impacted through either (i) zero inherent information yet a full perceptual range, or (ii) full inherent information yet zero perceptual range.

We illustrate the impact of this alternative representation under the chasm information field in Figure 16, where the background information is either low (Ω_0_ = 0.25), medium (Ω_0_ = 0.5) or high (Ω_0_ = 1). We observe that at higher background information levels, an individual is able to more efficiently navigate with a loss of perceptual range within an information void, compared to a reduced level of inherent information. A loss of perceptual range means that the individual will navigate based solely on its inherent information. Provided that there is a sufficient level of background information, navigation remains efficient. In contrast, for lower background information levels, navigation is more efficient with a loss of inherent information, compared to a loss of perceptual range. Here, the loss of inherent information is less significant, as the population relies on collective navigation to undertake efficient migration. These results suggest that different navigation strategies can be optimal, depending on the information available to the individual.

**Figure 16:**
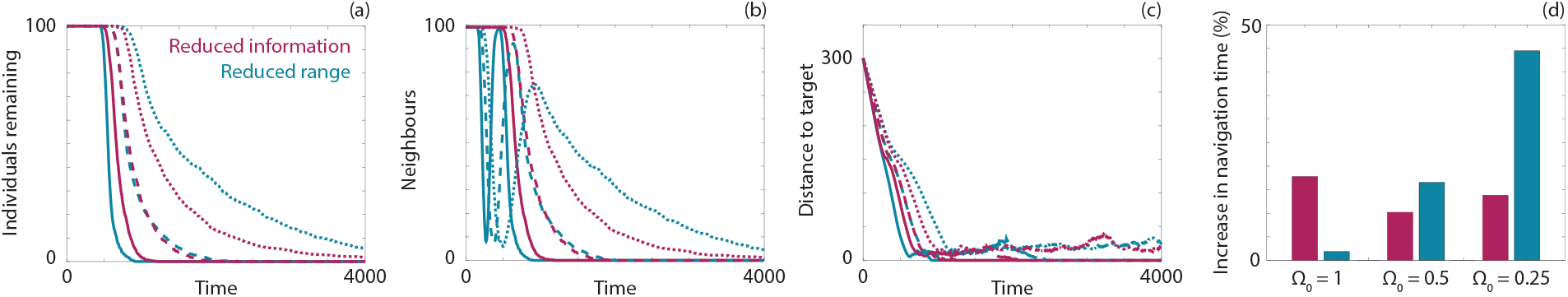
Navigation behaviour for 100 individuals in a chasm information field for (magenta) noise-dependent information and (blue) noise-dependent perceptual range. (a) The number of individuals remaining in the system over time. (b) The average number of neighbours within the perceptual range. (c) The distance between the target location and the average location of the individuals. (d) The increase in navigation time due to the presence of the information void, that is, relative to a constant information field with the same Ω_0_ value. Results are presented for Ω_0_ = 1 (solid), Ω_0_ = 0.5 (dashed), and Ω_0_ = 0.25 (dotted). Parameters used are *α* = *β* = 0.5, *r*_max_ = 500 and **c** = {**x** | 125 *≤* ||**x** *-***x**_target_|| *≤* 175}. All data are the average of ten realisations of the model.

### Fractional Brownian noise information fields

For further details on how to generate fractional Brownian noise information fields see [27]. We generate information fields, Ω_FBN_(**x**), according to process detailed in [27], and impose a scaling such that the noise is defined between zero and one. To generate distinct information-rich and information-poor regions, we transform the noise level according to obtain the information field:

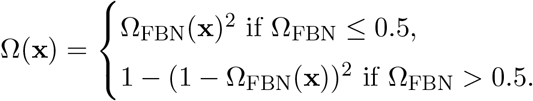

